# Biodiversity and local interaction complexity promote sustainable fisheries in large food webs

**DOI:** 10.1101/2022.12.08.519558

**Authors:** Alexandra Werner, Georg Albert, Ulrich Brose, Benoit Gauzens

## Abstract

On a global scale, fisheries harvest an estimated 96 million tonnes of fish biomass annually, making them one of the most important drivers of marine ecosystem biodiversity. Yet little is known about the interactions between fisheries and the dynamics of complex food webs in which the harvested species are embedded. We have developed a synthetic model that combines resource economics with complex food webs to examine the direct effects of fishing on exploited species and the indirect impact on other species in the same community. Our model analyses show that the sensitivity of the targeted species increases with its trophic level and decreases with its local interaction complexity (i.e. its number of interactions with prey, predators, and competitors). In addition, we also document a strikingly positive effect of community species richness on the resilience of the harvested species to this disturbance. The indirect effects on other species show specific patterns of spreading across trophic modules that differ systematically from how other disturbances spread across ecological networks. While these results call for further research on how human resource exploitation in general and fishery in particular affect ecological dynamics and biodiversity in naturally complex systems, they also allow for some cautious conclusions. Taken together, our results suggest that the sustainability concerning fishery yield and ecosystem integrity can be maximised by focusing the harvest on low trophic level species with a high local interaction complexity in high biodiversity ecosystems. In this sense, our complex network approach offers a promising avenue for integrating the necessities of generating economic revenue with the protection of natural biodiversity.

## Introduction

In the Anthropocene, humans are the dominant species on Earth (Moll et al., 2021), shaping ecosystems in various ways. One way is through resource consumption, where we have rerouted 20-40% of global net primary production to our benefit (Barnosky et al., 2012). In many food webs, these resource flows to humans are much stronger than any other trophic interaction, meaning humans can exert control and pressure on populations and ecosystems (Crabtree et al., 2019). One particularly well-studied and important type of human resource use is fishing, where we have caused trophic cascades (Darimont et al., 2015; Hoeks et al., 2020; Moll et al., 2021) regime shifts, and accelerated evolutionary pressures (Guerra et al., 2020). Despite this, *Homo sapiens* have traditionally been excluded in food web studies (Darimont et al., 2015) as we arguably differ from typical predator-prey behaviour, leading to limited theoretical knowledge on how human-caused disturbances spread through a network. This difference raises the fundamental question of how human exploitation links affect food-web dynamics and persistence.

With the advent of farming, humans decoupled from predator-prey dynamics and decreased prey populations (Darimont et al., 2015). As apex predators, we hunt or fish any sized animal we wish and seek rare species to maximise profitability (Courchamp et al., 2006). The most decisive factor on resource exploitation now is no longer physical or biological limitations but societal (i.e. quotas, moratoriums, market prices)(Ellis, 2015). We, therefore, have two very contrasting strategies for resource exploitation. Animal predators typically have density-dependent predation strategies, implying switching when prey species become scarce (Kondoh, 2006). Such behaviours facilitate coexistence among competing predators and add stability to food webs (Heckmann et al., 2012; Kondoh, 2006). However, harvesting by humans deviates substantially from such natural predation as it focuses on large (Darimont et al., 2015; Reynolds & Bruno, 2012) and rare species (Courchamp et al., 2006), mostly from higher trophic levels, which causes strong ecological effects (Estes et al., 2011). This difference suggests that we cannot understand the consequences of human resource exploitation by generalising predator-prey theory. Having played the leading role in prehistoric megafauna extinction (Hoeks et al., 2020; Muhly et al., 2013) and currently driving the ongoing biodiversity crisis (Mouillot et al., 2014), we must understand how humans, embedded in complex food webs, affect community stability and biodiversity.

No species lives in isolation, as every population, including humans, is embedded in a complex network that connects populations through their manifold trophic interactions and drives their response to a perturbation such as the biomass decrease of a predator. These complex interactions make it difficult to predict biomass outcomes in a food web after a disturbance. On the one hand, some features like omnivory (Fei & Kong, 2021) are expected to stabilise (reduce biomass change) food webs. On the other hand, it has been shown that the effects of disturbances (perturbations) in a food web decrease with increasing link or network distance from the affected species (Berlow et al., 2009), and that such disturbances are limited to closely interacting species within modules (Stouffer & Bascompte, 2011). The studies above obtained many of their results through simulations of species removal, which is a disturbance distinct from continuous resource exploitation such as human use. Therefore, we need to learn more about how disturbances caused by human resource exploitation spread in complex networks and how they would differ from those classically considered in the ecological literature.

Fisheries extract three to four times more biomass than any natural predator (Hansson et al., 2018) and exert direct pressures through harvesting and indirect pressures through trophic cascades that can alter community composition and species coexistence (Moll et al., 2021). Sustainable resource use thus goes beyond managing specific resource types at the population level. Instead, it must consider the complex organisation of ecological communities and the indirect effects that can emerge from it. By compiling the set of trophic interactions, food webs offer suitable tools to address the different facets a perturbation can have on ecosystems, from species extinctions (Stouffer & Bascompte, 2011) to provisions of ecosystem services (Keyes et al., 2021). To date, the modelling of fishery effects has been almost exclusively focused on system-specific studies of single food webs (such as Ecopath with Ecosim) (Pauly, 2000), simplified stage-structured consumer-resource models (Al-Darabsah & Yuan, 2018), or size-spectrum approaches capturing the biomass distribution across size classes (Blanchard et al., 2017). Multispecies size-spectrum models have provided insights into how different fishery management types affect biodiversity. Still, a more detailed understanding of how indirect effects on other species cascade through the communities can be gained through network approaches. Recently, a study linked the economic dynamics of fisheries to the ecological dynamics of complex food webs to show that harvesting high-biomass fish stocks, while initially rewarding, can undermine sustainability through indirect extinction cascades (Glaum et al., 2020). Similarly, it has been shown that different fishing strategies can trigger trophic cascades across complex food webs that amplify biomass fluctuations at lower trophic levels (Uusi-Heikkilä et al., 2021). However, detailed analyses of how human harvesting induces perturbation cascades through complex food webs have remained elusive.

Human behaviour, ecological processes, and institutional practices have been combined in social-ecological system research (Larkin, 2011). However, most are from an anthropocentric perspective (how humans are impacted), with rare theoretical investigations from an ecological view (Guerrero et al., 2018). Natural resource management plans are shifting from single-species management to integrative or ecosystem-type management plans (Howell et al., 2021). Still, the complexity and resources needed to create such models are barriers to policy uptake (Howell et al., 2021). Human resource exploitation differs from natural consumers as we value rarity (Courchamp et al., 2006). Therefore, understanding the effect of resource exploitation on ecosystems requires integrating economic resource models with complex ecological networks. In this context, we follow previous research on fisheries effects in complex food webs (Glaum et al., 2020; Uusi-Heikkilä et al., 2021) by investigating how such human-caused disturbances spread through a network. More specifically, we aim to study (1) how the response of harvested species to human pressure depends on its position and linkage patterns in the network and community biodiversity, (2) how this response propagates to other species through the network of trophic interactions and (3) whether this response depends on the type of industry harvesting the resources. We answer these questions by modelling the effect of two types of fisheries (small/large) on the biomass and coexistence of species in food webs.

## Method

### Model summary

The proposed model combines a food web generated from an adapted allometric trophic network model by Schneider et al. (Schneider et al., 2016) with simple economic drivers. We incorporate humans into the food web using a fishing fleet, removing targeted species biomass through fishing activities and dynamically adapting the fleets’ fishing intensity by changing vessel coverage or work intensity. Parameters and extended formulas are in **SI 1**.

### Food web topology

Our food webs contain three classes of species defined by the log_10_ body mass, *m_i_*, and metabolic type (plant, invertebrate, and fish). We generated food webs with 30 primary producer species (plant) and between 25 to 100 consumer species, with 2/3 of the latter set as invertebrates and the remainder as fish, *m_i_* is generated from a uniform distribution in [0,6] for producer species, [2,6] for invertebrates and [4,10] for fish (Schneider et al., 2016). The trophic niche of a consumer species (i.e. the body mass interval it feeds on) is defined by a hump-shaped Ricker function that quantifies consumer *i*’s probability (*L_ij_*) of successfully attacking and consuming prey *j*.

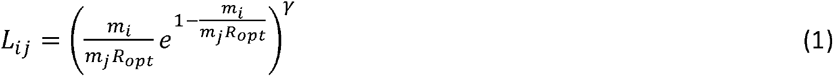

*m_i_* and *m_j_* are the body masses of the resource and consumer species, *R_opt_*, is the optimal predator-prey body mass ratio, and *γ* sets the width of the trophic niche. We assumed no interaction if *L <= 0.01*.

### Ecologic dynamics

We used a set of differential equations to determine the dynamics of nutrients *N_l_*, species biomass *B_i_* in our system. Plants compete for two nutrients of constant replenishment and lose biomass through herbivory and metabolism. Trophic relationships (predators and prey) and metabolism drive consumer biomass dynamics. The dynamics of nutrients are:

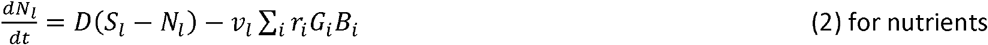

where *D* is the nutrient replenishment rate. The maximum nutrient concentration is *S_l_*, while *v_l_*, is the relative content in the plant species biomass. Producers grow due to their intrinsic growth rate *r_i_*, and specific growth factor *G_l_*,

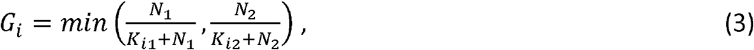

calculated from the nutrient concentration *Ni* and the nutrient uptake efficiency *K_i1_*, assigned for each species (Schneider et al., 2016). The producer and consumer dynamics are described through the following equations.

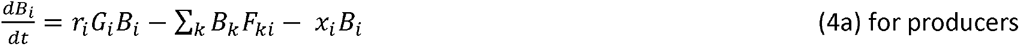

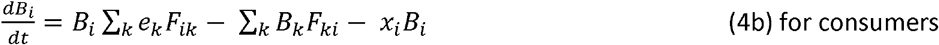

Where *k* represents consumer species and *F_ij_* describes the feeding rate of species *i* on resource *j*. Consumer growth scales with the assimilation efficiency *e_k_* which differs for plant and animal resource species (Lang et al., 2017). Decreases in biomass densities occur from consumption by consumers and metabolic demands 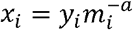, where *a* and *y_i_* differ for producers, invertebrates, and fish, respectively (Bland et al., 2019).

### Feeding rates

We calculated feeding rates *F_ij_* as

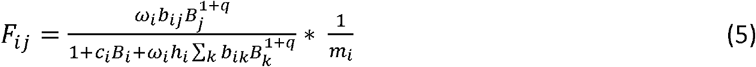

1+*q* is the Hill exponent, drawn for each food web (Schneider et al., 2016) and *c_i_* defines the consumer interference. The time a consumer spends attacking, consuming, and digesting a resource (handling time *h_i_*) depends on the body sizes of the consumer and resource scaled by the exponents *η_i_* and *η_j_*. It uses the scaling constant, *h*0, and is calculated as

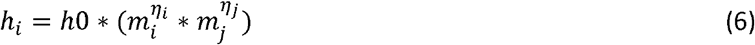

*ω_i_* is the relative consumption rate a predator has over its different prey, defined as one over the number of resource species of species *i*. We determine the encounter rate by the resource-specific capture coefficient, *b_ij_*, using an allometric relationship that differs depending on resource type *i* (herbivores and carnivores).

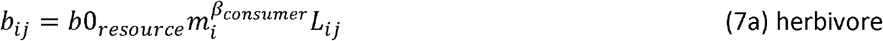

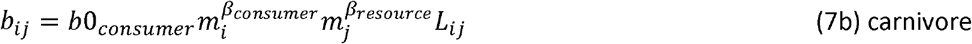

The encounter rate is only determined by the herbivore’s movement (**SI 2**), as plants do not move.

### Humans as a food web node

Humans affect the food web through the total amount of targeted fish species biomass harvested, *catch_VTotal_*, using the surface area covered by vessels in a fleet, *V_Total_*. We assume the fleet consists of identical vessels searching for and acquiring a specific species, equivalent to an open access system, with vessels entering or exiting depending on the profitability of the fishery (Bjørndal & Conrad, 1987). A fishery initially covers an area randomly drawn from [0.0001,0.001]. For the harvested species, **Equation 4b** becomes.

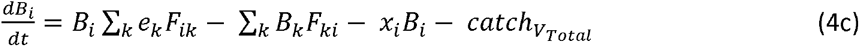

where

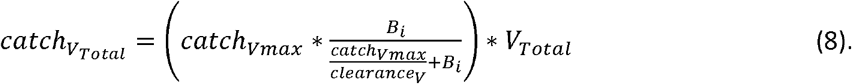

*catch_Vmax_* is the maximum biomass a vessel can catch (Ayunda et al., 2018), with *clearance_V_* being the surface area a vessel can clear of fish (a proxy for work). We derive *clearance_V_* in SI 3. The dynamics of *clearance_V_* and *V_Total_* depend on the difference between revenues made from harvesting and associated costs, *cost_V_*. The latter emanates from the necessity to uphold the ships’ operational readiness and comprises, *maintenance_V_*, and *clearance_V_*.

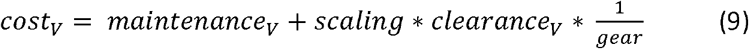

*scaling* controls the relative importance of *clearance_V_* to *maintenance_V_*, and *gear* is the type of equipment used by the vessel. Having better gear decreases *clearance_V_* cost.

Revenues depend on the price at which the market will buy a unit of harvested species *p_fish_*. They change according to the supply on the market (*catch_VTotal_*) and a base price associated with the harvested species (*p_base_*).

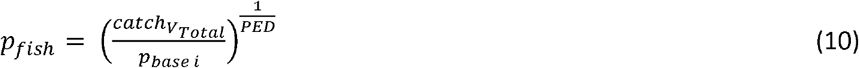

This formulation assumes a downward-sloping demand function (**SI 4**) where the elasticity of demand, *PED*, defines how sensitive the market price of harvested species is to a change in fish supply (Costello et al., 2016). We further assume larger species are valued higher by the market (Courchamp et al., 2006), adjusting *p_base_* to species *i* by multiplying it with the fraction of *m_i_* over the mean consumer body mass *m_n_* in the food web.

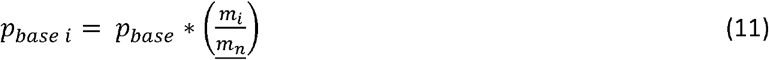

Selling *catch_VTotal_* generates fleet revenue (*revenue_V_Total__*).

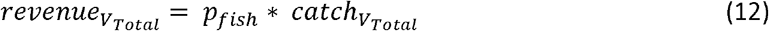

Subtracting the costs from the revenue gives the profit for the fishing fleet, *π_VTotal_*.

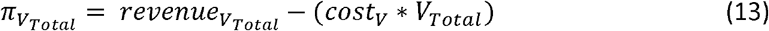

The number of vessels and the clearance change dynamically based on profit.

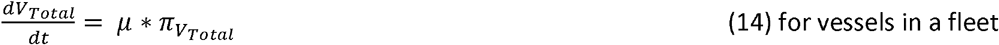

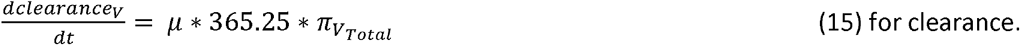

With positive (negative) profit, fleet and clearance increase (decrease). How quickly *π_VTotal_* translates to a change in *V_Total_* and *clearance_V_* is dependent on *μ*. As *clearance_V_* is associated with daily fishing activities, we multiply *μ* by the number of days in a year.

### Simulation set-up

We generated four food web groups of 25, 50, 75, and 100 initial animal species (**SI 5A**). These pristine food webs ran without harvesting for 100 years model equivalent to allow population stabilisation as a starting point for fishery simulations. A species became extinct when its biomass fell to 10^-6^. If less than 75% of fish species remained at the end of this first simulation, food webs were discarded. Food web generation continued until each food web group had 50 food webs. Final species biomass values from these non-harvested food webs are benchmarked against the final species biomass values from harvested simulations. We sequentially selected the species to be harvested over all consumer species in a food web. We then ran two models: one with the small fishery and one with the large fishery harvesting the selected species. These fisheries differ in clearance and gear used, with large fisheries being more efficient (**SI 6**). We repeated this setup for all food webs. In total, 200 food webs were used, consisting of 50 replicates for our four food web groups. We simulated the effect of harvesting on each animal species independently for each food web, leading to one simulation per animal species per food web (**SI 5B**). This was repeated for small and large fisheries, respectively.

To examine the impact of a fishing industry on a food web, we excluded data for harvested species that persisted after its fishing industry was not profitable enough to persist (density below 10^-6^). Some data points were considered outliers and removed from the analyses: when non-harvested species increased more than 100-fold in biomass at the end of the simulation run (1128 cases out of 2.6 million); and food webs where the harvested species had higher biomass at the end of the harvesting scenario (2 cases). If a data point was discarded for a given combination of complexity, harvested species and food web identity for the small scale scenario, we removed the result with the same complexity level, targeted species and food web in the large scale scenario.

### Descriptors

Using descriptors to determine the relationship between non-harvested and harvested species allows us to investigate how perturbations caused by harvesting spread through the food web and establish reasons for positive or negative biomass changes. *These are extrinsic characteristics that depend on the species harvested*. Using descriptors to identify the position of a species in the food web (trophic position) or how many network connections it has to other species (interactions) gives insight into the species’ characteristics. *These are intrinsic characteristics a species has independent of harvesting*. See **SI 7** for a list and description of the indicators used.

## Results

### Fishery scenario effects

We start our analysis by investigating whether the ecological consequence of changing the scale of fishing operations (small or large fisheries) differs between direct and indirect impacts for the harvested species and other species of the food web, respectively. We found that the large fishery always had a more significant impact on the harvested species and food web than the small fishery. More specifically, across all combinations of the other independent variables, the large fishery had roughly twice the effect on mean harvested species biomass (−31% vs −16%) and almost fourteen times the extinction rate of the harvested species as the small fishery (11.37% vs 0.82%). However, the patterns of how the direct effects on the harvested species and the indirect effects on the food web vary with our other independent variables are similar between both types of fishery. Despite these substantial differences between the two fishery types, we will thus present the following results for the large fishery scenario only, as the consequences of the small fishery are qualitatively consistent. Results of the small fishery can be found in **SI 9**.

### Fishery effect on the harvested species

We ran generalised additive models (GAMs) with predictors to explain harvested biomass change and harvested extinction. These predictors are the species richness of the food web and six characteristics of the harvested species (metabolic type, trophic level, number of prey species, number of predators, and number of competitors). Due to the strong correlation between body mass and trophic level in our generated food webs, we focused our analyses on trophic levels as they are comparable between food webs. We ran two independent GAMs for (1) harvested species biomass and (2) harvested species extinction probability with the five independent variables.

The best-ranked GAM (Based on AIC) for harvested biomass change used all six predictors to explain 92% of the variation. The best-ranked GAM for harvested species extinction probability explained 73% of the variation and only excluded metabolic type. We observed that species richness curbs biomass loss and decreases extinction risk (**Figure 1**). Therefore, harvested species lose proportionally less biomass and have a lower extinction probability the higher the species richness of the food web. Additionally, we find that vertebrates experience higher relative biomass loss when harvested than invertebrates (**Figure 1A**). This higher relative biomass loss leads to extinction probabilities of vertebrates that decrease with the species richness of the food web (**Figure 1B**).

**Figure 1.**
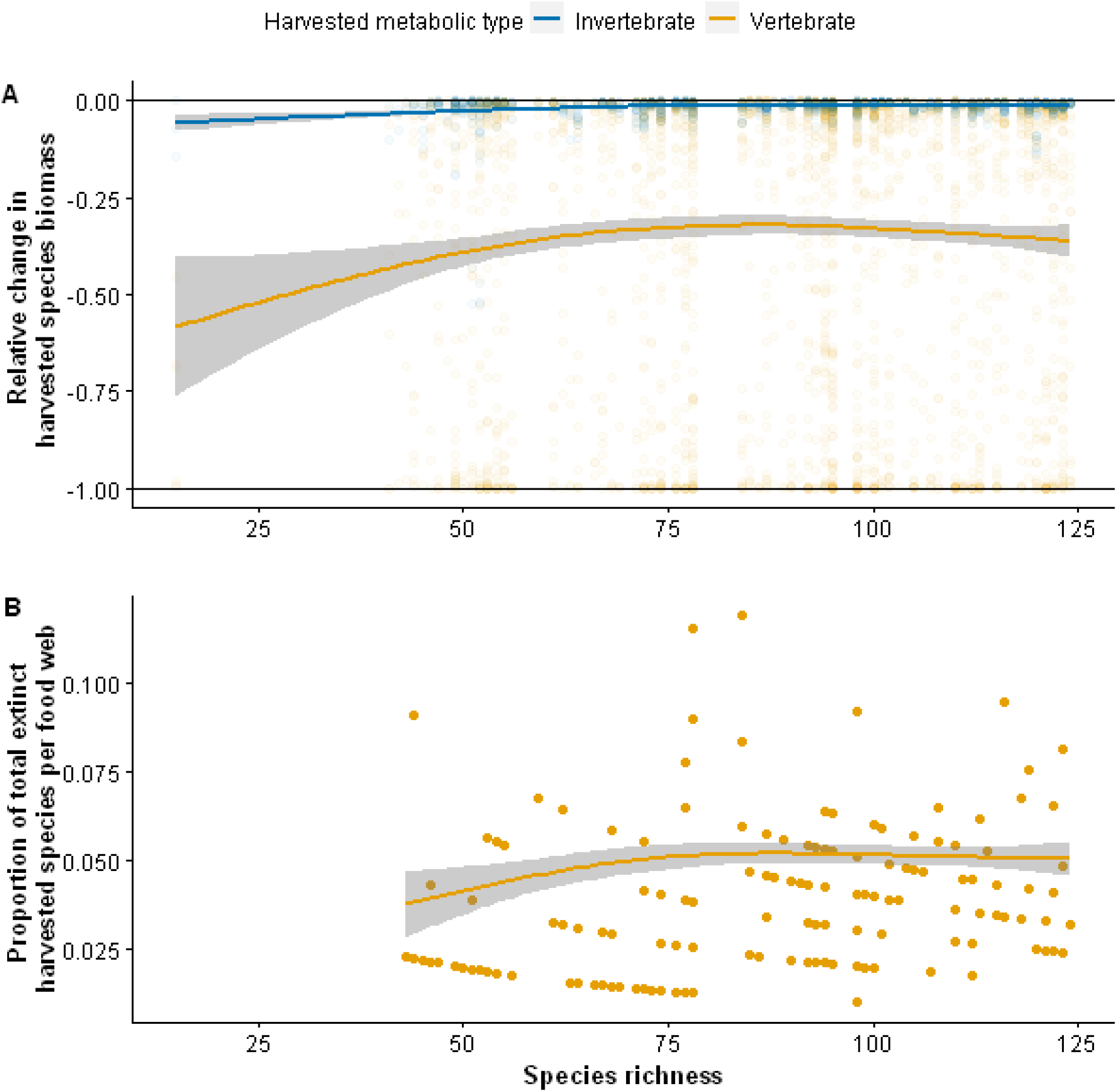
Relative biomass change and extinction proportion of the harvested species for a large fishery. (A) Relative biomass change for harvested species (y-axis) vs food web species richness. Thin black lines indicate no change (y = 0) and species extinction (y = −1). (B) The proportion of total harvested extinctions per food web (y-axis) vs food web species richness. The proportion is the sum of all extinct harvested species that occurred in that food web divided by the number of species at the start of the simulation. Lines represent GAM predictions and shaded area 95% confidence interval.

When harvested, species at higher trophic levels experience higher biomass losses than low-trophic level species (**Figure 2A**). While extinction counts are not limited to higher trophic positions (**Figure 2E**), they are almost exclusively restricted to species of trophic level four or higher. Generally, the fewer interactions a species has (fewer predators, prey species etc.), the higher the relative biomass loss (**Figure 2B-D**) and also the higher the extinction count (**Figure 2F-H**). Our results also indicate a limited overlap in the trophic levels occupied by both metabolic types, invertebrates and vertebrates (**Figure 2A**). The differences between the metabolic types emerging in our results could thus be an indirect effect of their trophic levels. To test for this possibility, we re-ran our analyses while restricting the data to trophic levels occupied by both metabolic types. These restricted GAMs show invertebrates lose less relative biomass than vertebrates (**SI 10** FigA), while the trends in biomass losses with the linkage patterns (number of predators, prey and competitors) are consistent between the two groups (**SI 11**, FigB-D). Together, these results suggest that relative biomass losses and extinction counts are (1) higher for vertebrates than for invertebrates, (2) increase with the trophic position, and (3) decrease with the number of interactions (with prey, predators and competitors) of the harvested species (**Figure 2**).

**Figure 2.**
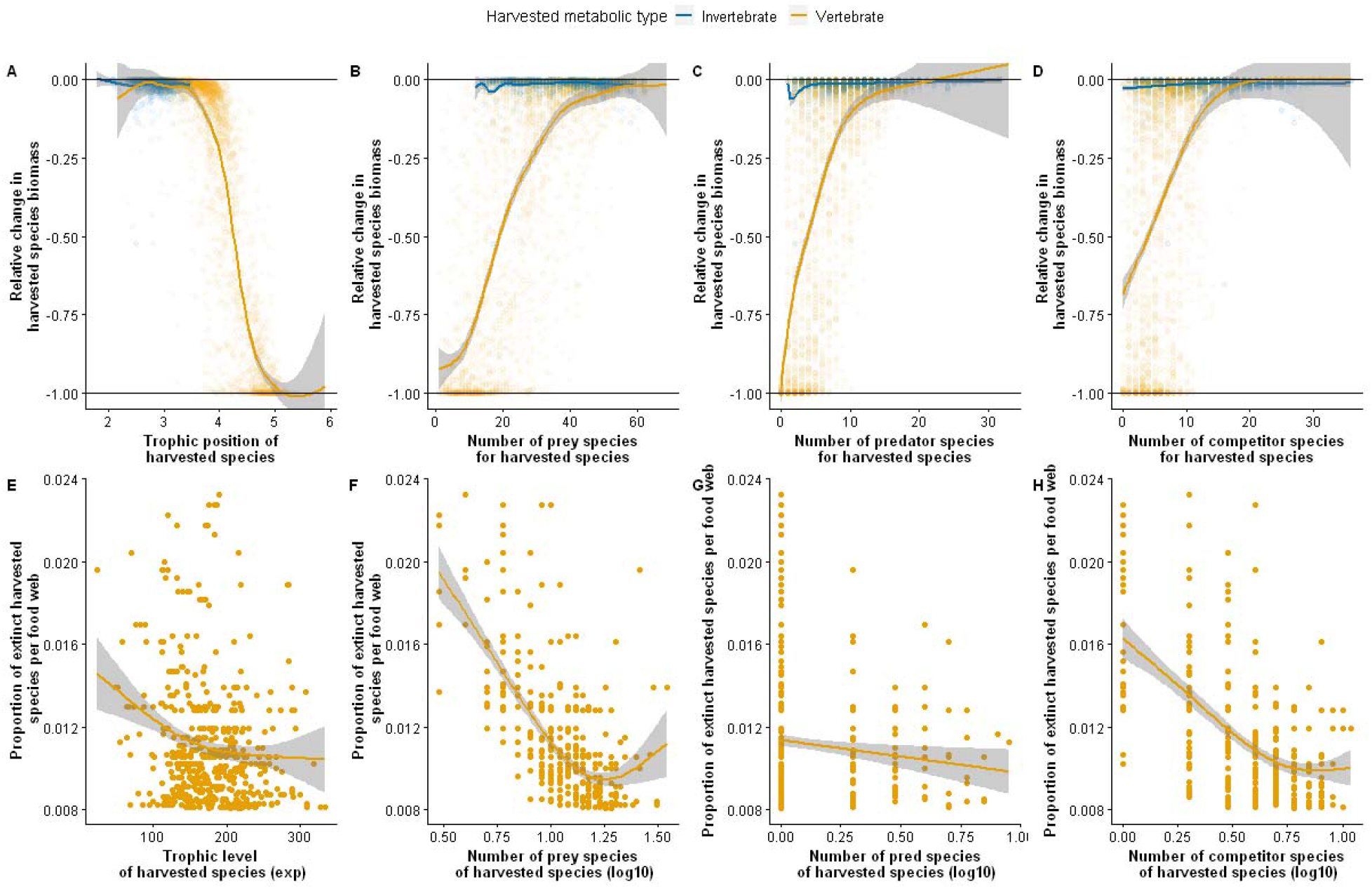
Total extinction counts and biomass changes in the harvested species for a large fishery. Relative change in harvested species biomass (y-axis) vs the harvested species trophic position (A), number of prey species (B), number of predators (C), and number of competitor species (D) - Thin black lines indicate no change (y = 0) and species extinction (y = −1). GAM with standard error for the relative change in harvested species biomass is calculated without extinct harvested species. The proportion of harvested species extinction per food web richness (y-axis) vs the harvested species trophic level (E), number of prey species (F), number of predators (G), and number of competitor species (H). No invertebrate went extinct. The proportion was calculated by dividing the extinct harvested species by the number of species in that food web

### Fishery effect on communities

In addition to the direct effects on the harvested species, fishing can also have indirect effects cascading through other species in the same food web. Here, we address how strongly non-harvested species’ relative biomasses are indirectly impacted by the extrinsic properties of modularity (same versus different module as the harvested species), link distance to the harvested species, respective role, and trophic position difference (difference in trophic level relative to the harvested species). First, we tested for differences in how the harvesting perturbation spreads depending on (1) whether a species is in the same or a different module of the food web and (2) the number of links distance to the harvested species. Second, we assigned roles for each non-harvested species (prey, competitor, predator, other-below, other-above) and plotted these against trophic position differences relative to the harvested species to see how the perturbation “wave” spreads in a food web.

Although the mean relative biomass of a food web, excluding harvested species, changes little (0.39%), some non-harvested species exhibited dramatic changes in relative biomass (increases up to 5635% and decreases to extinction-level). The spread (variance between species in a food web) of relative biomass change is tenfold greater within the same feeding module than in a different feeding module. This variance decreases across the entire gradient of link distance by three orders of magnitude with increasing link distance from the harvested species (**Figure 3A**).

**Figure 3.**
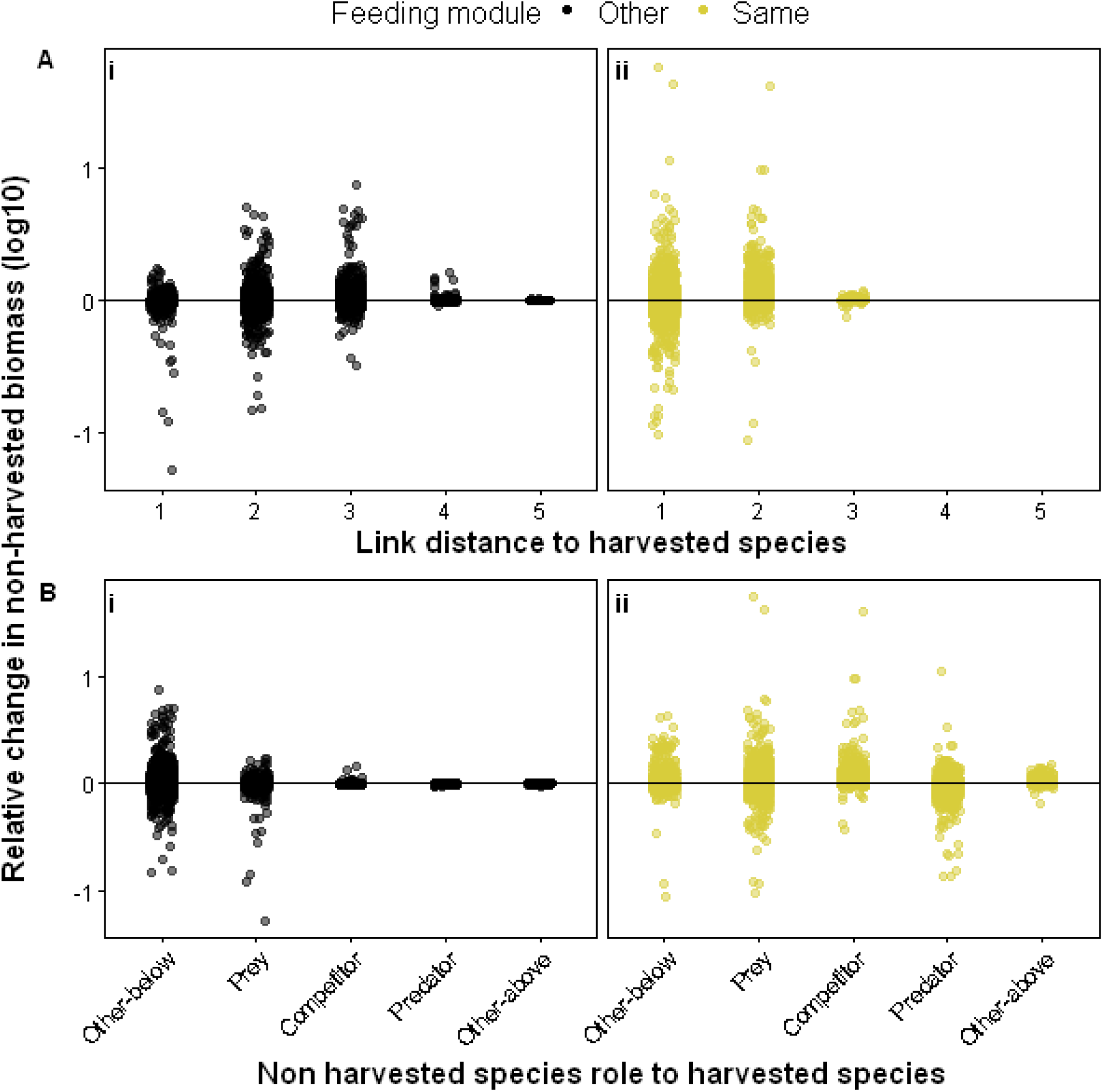
Relative biomass change in non-harvested species for all food webs per link distance to, and role for, the harvested species, large fishery. The relative change in non-harvested species biomass (y-axis) is the harvested biomass divided by the pristine biomass for each non-harvested species. Link distance (x-axis) is the trophic position of the non-harvested species subtracted by the harvested species’ trophic position. Roles are assigned based on network connection and Jaccard Index. Species that interact closely are assigned to the same module. For each plot, the horizontal line indicates no change (y = 0). Figure 2A Large Fishery results showing link distance and feeding module; Figure 2B Large Fishery results showing role and feeding module. Extinct species have been removed.

In a different feeding module, the spread of the perturbation is reduced almost six times across the entire link range (five links distance to the harvested species), but within link distances, the variance first increases between link distances 1, 2 and 3 before decreasing (**Figure 3Ai**). This contrast illustrates a difference in how the perturbation decays with increasing link distance when comparing non-harvested species in the same versus another module. Within the same module, we found a systematic decay in the biomass effects of this perturbation with link distance to the harvested species. In other modules, our results reveal a hump-shaped pattern of biomass changes. Subsequently, we compared the biomass changes across the different roles of the non-harvested species. In other modules, we systematically found weaker effects on the harvested species’ competitors, predators, and Other-above than their prey and Other-below (**Figure 3Bi**). In the same modules, we find similar variances in biomass changes across the roles of the predator, prey, competitor and Other-below, and only Other-above exhibits a more narrowly constrained range of weaker responses to the perturbation. Together, these results reveal that the perturbation spreads similarly to all species roles in the same module, except for Other-above, while systematically decaying with increasing link distance to the harvested species. However, in other modules, the perturbation mainly spreads to prey and Other-below. The decay in perturbation effect follows a hump-shaped relationship with link distance to the harvested species.

Combining modularity, link distance, and non-harvested species role (**Figure 4**) allows us to see which roles react with biomass change at each link distance in the food web. In a different module, the variance is first driven by prey species at link distance one and then by other-below species with link distances two and three (**Figure 4A**). Unusually, the widest biomass change spread is seen for other-below species, not directly associated with the harvested species. In the same module, the largest variations are seen first in prey and predator species at link distance one and then by other-below and competitor species at two link distances from the harvested species (**Figure 4B**).

**Figure 4.**
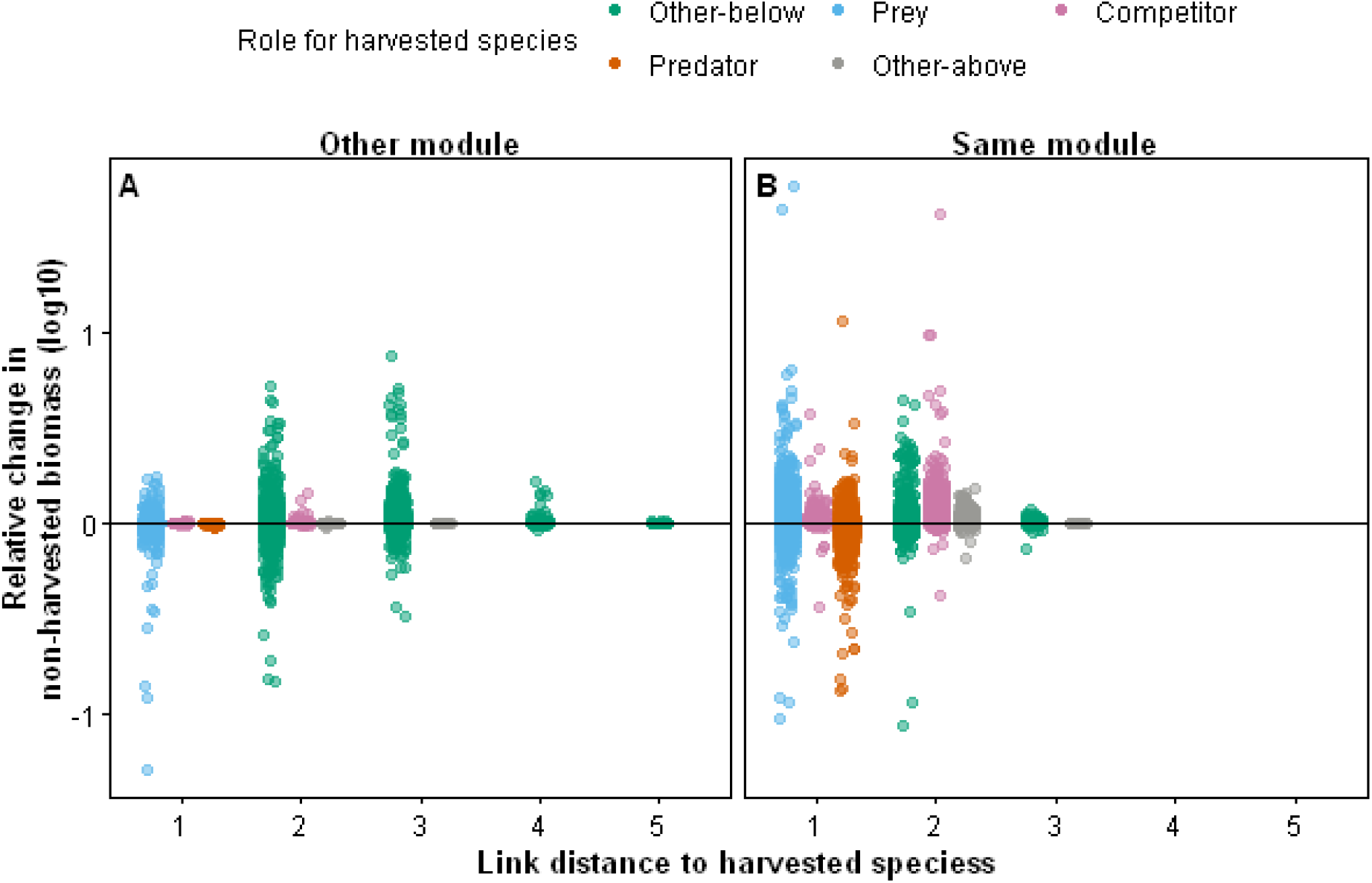
Relative biomass change in non-harvested species per role and link distance to harvested species for all simulated food webs, large fishery. Relative change in harvested species biomass on a log10 scale (y-axis) versus the non-harvested species links distance to the harvested species (x-axis). Thin black lines indicate no change (y = 0). Figure 3A Large Fishery results from a different feeding module; Figure 3B Large Fishery results from the same feeding module.

We then plot the data from **Figure 4** against trophic position differences to examine how the non-harvested species’ biomass changes depending on its distance from the harvested species. **Figure 5** visualises how a human-induced disturbance spreads through a randomly chosen example of our simulated food webs. In the same module, the disturbance originates from a trophic position difference of 0 (**Figure 5B**). It spreads with decreasing amplitude through lower trophic positions and dissipates entirely at higher trophic positions (trophic position difference greater than 1.5). Competitors profit, as they have the greatest mean biomass change, which is almost twice the mean for preys, which also have a positive mean despite the competitor’s biomass gain. Predators are significantly disadvantaged, having the only negative mean biomass change from all other roles.

**Figure 5.**
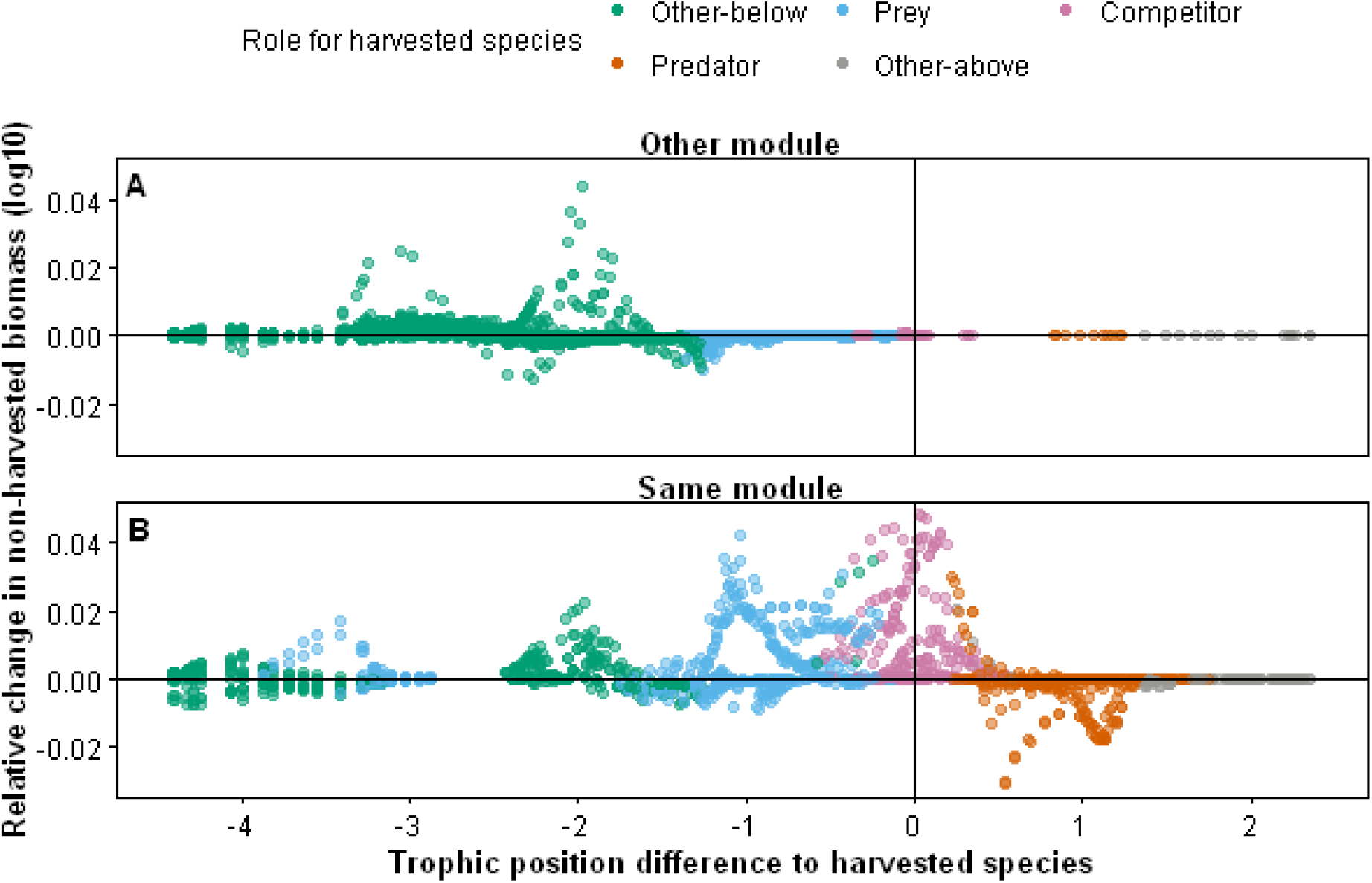
Relative change in non-harvested biomass for a random food web. The relative change in non-harvested species biomass (y-axis) is the harvested biomass divided by the pristine biomass for each non-harvested species. Trophic position difference (x-axis) is the trophic position of the non-harvested species subtracted by the harvested species’ trophic position. For each plot, the horizontal line indicates no change (y = 0). Figure 4A Large Fishery results from a different feeding module; Figure 4B Large Fishery results from the same feeding module.

Based on the cascading pattern from right to left, perturbations spread from the same feeding module (**Figure 5B**) into other feeding modules exclusively via lower trophic positions (−1 trophic difference and lower) (**Figure 5A**). Minimal variation occurs in a different module higher than the −1 trophic position difference. Together, this suggests top-down control in our food webs. Other-below has the largest increase in mean biomass change, more than three times higher than competitors. The mean biomass change for prey decreased the most. Together, these results show a systematic pattern of harvest-perturbations mainly affecting the harvested species’ prey, predators, and competitors in the same module, while Other-below spreads to other modules.

## Discussion

Following recent work (Glaum et al., 2020; Uusi-Heikkilä et al., 2021), we generated a model integrating the population dynamics of complex food webs with the economics of a fishery. By explicitly including humans in complex food webs, we addressed how the effects of resource exploitation cascade through food webs depending on the species’ intrinsic and extrinsic properties. Confirming prior studies, we find large fisheries impose a stronger impact on the food webs than a small-scale fishery, but the effects are the same qualitatively. We find that a species’ harvest sensitivity is determined by their position in food webs, local interaction complexity, and metabolic type. Interestingly, our simulations of complex systems reveal that biodiversity buffers the effects of the fishery as the harvested species lose less biomass and experience lower extinction probabilities in species-rich food webs.

### Fishery effects in complex systems

Overall, our simulated large fishery had a greater impact on biomass and extinction rates than the small fishery, initiating strong top-down effects. These results are qualitatively similar to what is expected from ecological theory (Oksanen et al., 1981) and real-world results (Crilly & Esteban, 2013; Guyader et al., 2013).

### Harvested species – Direct effects

Food-web theory and empirical observations show that high-trophic level species are generally most prone to suffer from direct and indirect effects of perturbations (Binzer et al., 2011; Reynolds & Bruno, 2012). Corroborating these findings, we observed similar trends in our results, with species occupying higher trophic positions being more affected by harvesting (higher number of extinctions, stronger biomass loss). An explanation for this sensitivity to disturbances is found in the lower biomass density of species from higher trophic levels. The depletion of energy between predators and prey (assimilation efficiencies and species metabolic rates) accumulates through trophic levels, strongly constraining the biomass level of species at high trophic levels (Binzer et al., 2011). Therefore the sensitivity of high trophic levels to disturbances can thus be found in the food-web energetics that constrains their biomass densities.

Interestingly, our simulations of complex food webs also reveal the novel finding that the sensitivity to being harvested decreases with the local interaction complexity (the number of interactions with prey, predators and competitors). High local interaction complexity allows harvested species to switch resources depending on their relative abundances (Zhao et al., 2019) and have a low niche overlap with competitors (Allesina & Levine, 2011; Capitán et al., 2015), promoting their persistence. This argues for the importance of preserving the resource-harvested species and their set of interactors.

Therefore, our results support the conservationist view that less-connected species (often specialists) are the most sensitive to harvest and that well-connected species (generalists) already favoured in disturbed environments gain a competitive advantage in ecosystems under high harvest pressure. This sensitivity increases with the trophic level of the harvested species. We found sensitivity to also be higher for vertebrates than invertebrates because of their higher per unit metabolic rate, increasing their energetic expenditures and reducing their biomass densities.

Additionally, we found that the species richness of the food web buffers the negative effect of harvesting. We argue that species richness boosts the potential for network connections and, thus, the number of energy pathways that help harvested species compensate for disturbances. These novel results imply that preserving local biodiversity is then in the economic interest of fishery managers as it buffers changes in biomass density and increases the long-term sustainable yield.

### Non-harvested species – Indirect effects

Network theory and empirical studies have shown that perturbations induced by species removal are contained in the same feeding module as the harvested or disturbed species (Stouffer & Bascompte, 2011) and decrease with increasing link distance from the harvested species (Berlow et al., 2009). At the same time, some studies find perturbations affecting species from all modules in the specific case of fisheries (Pérez-Matus et al., 2017). Our results agree with both when we look at modularity, non-harvested species’ role, and link distance, together.

We observe that the magnitude of the disturbance is the largest in the harvested species’ module and immediate network neighbourhood (the harvested species’ prey, predators, and competitors). However, complex feedback loops must occur as the perturbation spreads to other modules at farther link distances. For species occurring in a different module than the harvested species, the perturbation is not the strongest for predators or prey species (one link distance) but for species at two or more link distances, usually at two trophic levels below the harvested species. Therefore, this suggests weak perturbations can spread through the network by different trophic routes and strongly affect very distant species. These findings do not match theory predictions derived from simplified food web representations or species removal simulations (i.e. mimicking extinctions), highlighting the importance of addressing mechanisms of how harvesting as a press perturbation causes chains of indirect effects in complex food webs.

Another deviation from theoretical predictions is the meagre biomass response of species from the highest trophic levels that are not consumers of the harvested species. Simulations and empirical evidence show top predators are the most vulnerable to extinction (Eklof & Ebenman, 2006). Such little biomass change in our food-web simulations can be explained by their ability to forage resources from different modules (Guimarães, 2020) simultaneously. Connecting modules allows them to compensate for energy flow disturbances at lower trophic levels in one module by shifting their diet to similarly-sized species to the harvested species but from other modules that exhibit almost no variation. Therefore, the modularity structure of food webs may have a dampening effect favouring the persistence of top species when perturbations only occur locally.

Our study did not find a systematic classic trophic cascade, as some species were unaffected or responded oppositely to what was expected. Indeed some prey species in our food webs decreased in biomass. Griffen et al. found that predator richness generally did not suppress prey more than its most effective predator species (Griffin et al., 2013). Higher-order emergent effects are likely occurring, where it is unknown if removing a species will increase its prey population based on how its competitor species interact with it and each other (Tekin et al., 2018) or through long feedback loops (Gauzens et al., 2016). Fisheries can therefore initiate indirect effects, and their control of the targeted species can free an ecological niche for some competitors that can reach higher biomasses. This indirect effect would correspond to an “indirect mutualism” that fisheries have on the competitors of its target species.

Taken together, our results on the sensitivity of the food webs’ top species to harvesting deviates substantially from general ecological disturbance theory, and the blurring of trophic cascades in complex food webs highlights the importance of studying harvesting disturbances in complex systems to come to a predictive understanding of human impacts on natural biodiversity. Further work may explain why disturbances enter other modules via lower trophic levels and help capture some of the complex feedback loops occurring within and between modules. Our results demonstrate that combining fisheries models with complex food webs opens the possibility of considering trophic cascades and indirect effects that can cause feedback loops on the harvested species. Therefore, our models build a foundation for more holistic ecosystem-based fisheries management by going beyond the impact of individual exploited species.

### Future Directions

In nature, no species lives in isolation, and our model approach makes a step toward incorporating ecological complexity in the study of resource harvesting by economic actors. However, ecological complexity has many facets, and like any modelling study, our approach included one of these dimensions, whereas others remained simplified. One of these simplified dimensions is the age- or stage-structure of the fish populations. Such structured population models, where the biomass of a species is distributed across various stages and body sizes (Bland et al., 2019), are increasingly used in fishery management. Similarly, size-spectrum models have also provided insightful results on how fisheries affect biomass distributions across species and size classes, leading to clear recommendations on mitigating biodiversity loss (Blanchard et al., 2014, 2017). Despite their increased use, these modelling approaches also ignore a dimension of ecological complexity, the surrounding food webs, which has been the focus of our study. Although there has been little research on integrating structured populations or size-spectra with complex food webs (Blanchard, 2011), this could be a fruitful avenue for future research on fisheries impacts.

Our approach considers a single fishery impacting a unique target species. While this approach helps better apprehend the consequences of harvesting and the interconnection between social-economic and ecological network dynamics, more factors might be important to test. Single-species fisheries are rare and do not exist in isolation, as other fisheries can catch the predator or prey of that species. This is a major criticism of single-species management (Hilborn, 2011). The dynamics of our results would change if more than one perturbation were occurring, say through multiple fisheries. Similarly, by-catch, a widespread phenomenon, also impacts population dynamics and could amplify the effects shown in our study. Its effect, however, could also dampen the strong effects seen at higher trophic levels as the fishery starts to act as a top predator, impacting the competitors of the harvested species. This suggests that multispecies fisheries and by-catch consequences represent important future additions to our network approach.

### Conclusions

By linking theoretical food web ecology and socio-economic approaches, we have expanded our understanding of how a human-caused disturbance by harvesting one species moves through a complex system of trophic interactions in ecosystems. Following up on recent work combining economic fishery models with complex food-web dynamics (Glaum et al., 2020; Uusi-Heikkilä et al., 2021), our results provide a characteristic risk profile for an intensive fishery, suggesting that the intrinsic characteristics of a harvested species (species richness, trophic level, metabolic type, and the local interaction complexity) strongly influence its sensitivity. In a network context, these characteristics can be integrated by the generic explanation that the more energy pathways surround a species, the more resilient this species will be to being harvested. Remarkably, our results also show that community species richness mitigates the negative effects of intensive fishing on exploited species. In addition, our dynamic network approach also revealed differences between human harvesting and other types of disturbances regarding cascading effects on the species community. Among the differences from classical network perturbation theory predictions, are the complex feedback within and between modules. They highlight that harvesting is a special type of disturbance. It is a press disturbance that increases in strength as the biomass density of harvested species decreases because prices in economic markets favour rare resources. This creates a clear separation from other types of disturbance, such as species extinction (single disturbance (Curtsdotter et al., 2011), global change stressors such as warming (press disturbance affecting all species (Binzer et al., 2016) irrespective of their density), and predation (e.g., by invasive species, positive density dependence of disturbance severity (David et al., 2017)). This advocates for further research on how fisheries and different types of harvesting cause disturbance that spreads through complex ecological networks. Using dynamic networks, our model produced results that indicate how to increase the sustainability of fishery yields while protecting the integrity of surrounding food webs. Namely, we found that sensitivity to overfishing decreases with the local interaction complexity and increases with the trophic level of species harvested. Moreover, our results argue for biodiversity protection to increase the sustainability of fisheries yields. We anticipate that such an in-depth look at the complexity of natural ecosystems will help provide clues for conserving biodiversity while maximising sustainable yields through ecosystem-based fisheries management.

## Supporting information

Supplementary information for the manuscript.

## Acknowledgements

AW, GA, UB, BG gratefully acknowledge the support of iDiv funded by the German Research Foundation (DFG–FZT 118, 202548816).

## Conflict of Interest

The authors have no conflict of interest to declare

## Author Contributions

Alexandra S Werner: conceptualization (equal), data curation (equal), formal Analysis (equal), investigation (lead), methodology (equal), visualization (equal), writing – original draft (lead), writing – review & editing (equal). Ulrich Brose: conceptualization (equal), funding acquisition (lead), methodology (equal), writing – review & editing (equal). Georg Albert: conceptualization (supporting), visualization (supporting), writing – review & editing (equal). Benoit Gauzens: conceptualization (equal), data curation (equal), formal Analysis (equal), methodology (equal), software (Lead), writing – review & editing (equal)

## Data availability

no data used

## Notes

### Competing Interest Statement

The authors have declared no competing interest.

