## Supplementary information for the manuscript. for "Biodiversity and local interaction complexity promote sustainable fisheries in large food webs"

## SI 1

### Table 1 Parameter symbols, meaning, values, and units

| Symbol in paper | Julia Computer Code Symbol | Meaning | unit | value | Reference |
| --- | --- | --- | --- | --- | --- |
| Ecological Parameters | | | | |  |
| Setting up | | | | |  |
| subscripts *_i j k_* | pred, prey | Consumer relationship between species _i,j,k_ | - | - |  |
|  | P InV V A | Plant Invertebrate  Vertebrate Animal | - | - |  |
| *m* | bm | Log _10_ body mass | g | Uniform dist.  μ_P_ = (10^0^, 10^6^)  μ_InV_ = (10^2^, 10^6^)  μ_V_ = (10^4^, 10^10^) | Schneider et al., 2016,  modified |
| *B* | bioS | Biomass density of plant, animal | g*m^-2^ | calculated |  |
|  | num_nutrient | Number of nutrients | - | 2 | Schneider et al., 2016 |
|  | num_plant | Number of plant species | - | 30 | Schneider et al., 2016 |
|  | num_invertebrate | Number of invertebrate species | - | 17 - 67 |  |
|  | num_vertebrate | Number of vertebrate species | - | 8 - 33 |  |
|  | num_vessel_types | Number of vessel types | - | 1 |  |
| Food web structure | | | | |  |
| *L* | L | Establishes potential links between species | - | calculated |  |
|  | foodWeb | Establishes probability of links existing | - | calculated |  |
| *R_opt_* | Ropt | Optimal consumer-resource body-mass ratio | - | 100 | Schneider et al., 2016 |
| *γ* | γ | Ricker function width | - | 2 | Schneider et al., 2016 |
| *e* | exp | natural logarithm base |  | 2.7182…. | - |
| Feeding rates | | | | |  |
| *F_ij_* | FF | Feeding rate or feeding function |  | calculated |  |
| *ω* | ω | Relative consumption rate  (1/# resources for species *i*) |  | calculated |  |
| *q* | q | Hill exponent | - | Norm. dist., μ = 0.5, σ = 0.2 | Schneider et al., 2016 |
| *c* | c | Consumer interference | - | Norm. dist., μ = 0.8, σ = 0.2 | Schneider et al., 2016 |
| *h* | h | Handling time | g*s^-1^ | calculated |  |
| *h_0_* | h0 | Scaling constant |  | 0.4 | Schneider et al., 2016 |
| *B_0_* | b0 | Capture coefficient; half-saturation density |  | Herbivores: 3500.0  Carnivores: 15.0 | Ryser et al., 2019 |
| β | β | Power-law relationship scaling |  | Herbivores: 0.19  Carnivores: 0.42 | Hirt et al., 2017 |
| *η* | η | Scaling constant |  | Norm. dist.  ηprey : μ = -0.48, σ = 0.03  ηpred: μ = -0.66, σ = 0.02 | Schneider et al., 2016 |
| Population dynamics | | | | |  |
| *e_k_* | ae_P , ae_A | Assimilation efficiency; conversion efficiency |  | P: 0.545  A: 0.906 | Lang et al., 2017 |
| *x_i_* | X | Metabolic demand |  | calculated |  |
| *x_i_* | x_P , x_InV , x_V | Metabolic rate |  | P: 0.138  Inv: 0.314  V: 0.88 | Schneider et al., 2016  Schneider et al., 2016  Bland et al., 2019 |
| *a* | - | Metabolic rate scaling constant |  | P: 0.25  Inv: 0.15  V: 0.11 | Schneider et al., 2016  Bland et al., 2019  Bland et al., 2019 |
| *r* | r | Intrinsic plant species growth rate | g*s^-1^ | r_i_ = m_i_^-0.25^ |  |
| *G* | CalcG | Plant species specific growth factor |  | Calculated, see Schneider et al., 2016 |  |
| *N* | bioS[num_nutrient] | Nutrient concentration | g*m^-2^ |  |  |
| *K* | K | Half saturation density of nutrient; nutrient uptake efficiency |  | Uniform dist.  0.1-0.2 | Schneider et al., 2016 |
| *D* | D | Global nutrient turn over rate; replenishment rate |  | 0.25 | Schneider et al., 2016 |
| *S* | S | Nutrient supply concentration |  | Norm. dist.  μ = 100, σ = 2 |  |
| *v* | v | Relative nutrient content in plant species biomass |  | v_1_ = 1  v_2_ = 0.5 | Schneider et al., 2016 |
|  |  | Extinction threshold |  | 0.000001 | Schneider et al., 2016 |
| Economic parameters | | | | |  |
| *catch_Vmax_* | catch_max | Maximum fish a vessel can catch per second | g*s^-1^ | 23 | Ayunda et al., 2018 |
| *μ* | μ | Economic flexibility | 1/m^-2^ * €^-1^ | 0.1-1.0 |  |
| *V_speed_* | V_speed | average vessel speed | m/s | 1.5 | FAO, 1980 |
| *Net_width_* | fish_net | fishing net width | m | 3.0 | FAO, 1980 |
| *fishing hours* | fish_hour | Hours per day spent fishing (fraction) | - | 0.58 | ILO, 2004 |
| *active fishing* | active_fishing | Fishing hours spent actively fishing (fraction) | - | 0.5 |  |
| *fishing days_max_* | fish_day_max | Maximum days per year spent fishing (fraction) | - | 0.69 | Guiet et al., 2019 |
| *fishing days_min_* | fish_day_min | Minimum days per year spent fishing (fraction) | - | 0.69 | Guiet et al., 2019 |
| *clearance_Vmax_* | V_clearance_max | Maximum vessel clearance rate | m^2^*s^-1^ | calculated |  |
| *clearance_Vmin_* | V_clearance_min | Minimum vessel clearance rate | m^2^*s^-1^ | calculated |  |
| *clearance_V_* | V_clearance | Realised vessel clearance rate | m^2^*s^-1^ | calculated |  |
| *B_0V_* | b0V | Vessel half saturation density | g*m^-2^ | calculated |  |
| $PED$ | elasticity | how sensitive the market price of fish is to a change in fish supply | - | -1.15 | Costello et al., 2016 |
| *p_Base_* | γ_fish | fish price coefficient | €*g^-1^ | 0.00129 | SEAFDEC, 2015 |
| *scaling* | cost_scale | Attack cost scale | €*m^-2^ | 5-100 |  |
| *maintenance_V_* | maintenance | cost of maintaining a vessel ready to leave port | €*s^-1^ | 0.05-0.1 |  |
| *time_Handling_* | - | Handling time | s*g^-1^ | - |  |
| *β_V_* | β_V | Scaling exponent for attack rate | - | 2.0 |  |
| *revenue* | revenue | Price consumers are willing to pay based on supply | €*s^-1^ | calculated |  |
| *cost_VTotal_* | cost_total | Total cost of fleet | €*s^-1^ | calculated |  |
| *cost_V_* | cost | Cost of one vessel | €*s^-1^ | calculated |  |
| $\pi_{V_{Total}}$ | profit | Positive or negative profit based on whether demand price covers cost of fleet | €*s^-1^ | calculated |  |
| *V_Total_* | bioS[(num_nutrient+num_plant+  num_animal+num_vessel_types)] | Area covered by vessel or fleet | 1/*m^-2^ | calculated |  |
| ${catch}_{V_{Total}}$ |  | Total biomass caught | g*s^-1^ | calculated |  |
| $V_{Total}$ |  | Total area covered by the fleet | 1*m^-2^ | calculated |  |
| ${catch}_{Vmax}$ |  | maximum biomass a vessel can catch | g*s^-1^ | calculated |  |
| ${clearance}_{V}$ |  | the surface area a vessel can clear of fish | m^2^*s^-1^ | initial  0.6967657933852107 |  |
| $gear$ |  | vessel equipment: net size and vessel speed | m^2^*s^-1^ |  |  |
| $p_{fish}$ | fish_price | price the market will buy fish at | €*g^-1^ |  |  |
| ${revenue}_{V_{Total}}$ | revenue | Selling all caught fish | €*s^-1^ |  |  |
| $\mu$ | $\mu$ | How quickly the number of vessels or clearance changes based on profit |  |  |  |

## SI 2

### Expanded Method formulae and explanations

The formulae 7a and 7b differ as encounters depend on the movement of consumers and resources (capture coefficient, *b0_resource_* = 3500.0, *b0_consumer_* = 15.0 (Ryser et al. 2019), scaling allometrically with the power-law relationship scaling exponent, β*_resource_* = 0.19, β*_consumer_* = 0.42 (Hirt et al. 2017)).

Hirt, M. R. et al. 2017. The little things that run: a general scaling of invertebrate exploratory speed with body mass. - Ecology 98: 2751–2757.

Ryser, R. et al. 2019. The biggest losers: habitat isolation deconstructs complex food webs from top to bottom. 286: 8.

## SI 3

### Initial clearance

0.6967657933852107, is based on the intercept of a Type II functional response (**Equation A**) and its inverse (**Equation B**), representing a choice in strategy to focus on abundant and less valuable prey (↓ costs ↓ pay off) or focusing on rarer but more valuable prey (↑ costs ↑ pay off). We expect an adaptive clearance rate to stay within the limits of these two equations. The intercept is the starting clearance value for every vessel.

A.	${clearance}_{a} =({clearance}_{Vmax}- {clearance}_{Vmin})*\frac{{B_{i}}^{\beta_{V}}}{{B_{0V}}^{\beta_{V}}+ {B_{i}}^{\beta_{V}}}+ {clearance}_{Vmin}$	B.	${clearance}_{V_{b}} = \frac{1}{\frac{1}{{clearance}_{Vmax}+ {clearance}_{Vmin}}*\frac{{B_{i}}^{\beta_{V}}}{{B_{0V}}^{\beta_{V}}+ {B_{i}}^{\beta_{V}}}+ \frac{1}{{clearance}_{Vmax}}}$

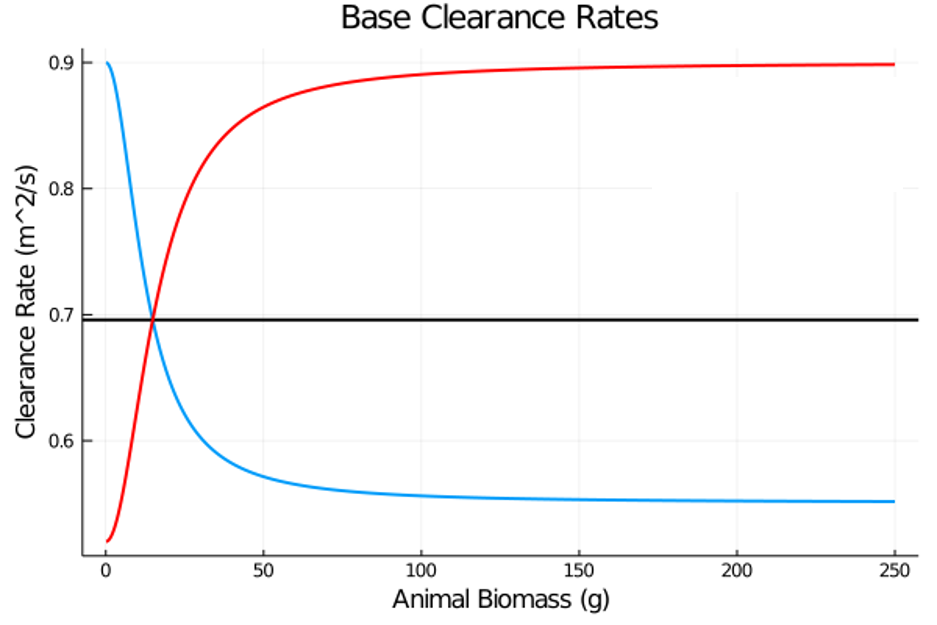

*Clearance types and their relation to animal biomass, used in our study: equation a - red, equation b - blue, intercept - black*

## SI 4

### Downward sloping demand function

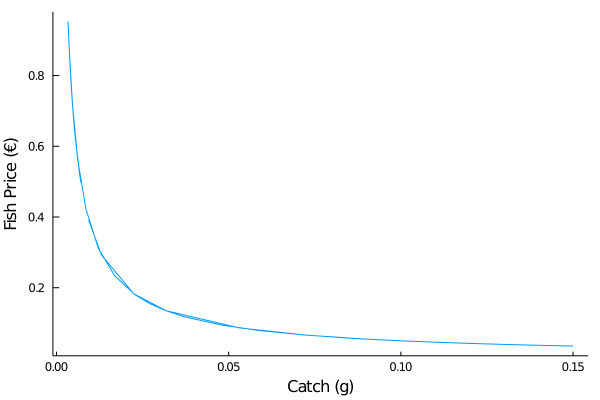

*How the price a fish is sold for changes with supply on the market. The Fish Price (y-axis) is the value the market is willing to pay for a gram of species i catch (x-axis). More fish on the market garners a lower market price.*

## SI 5A

### Summary of food web sizes and metabolic types

| **Initial Animals** | 25 | 50 | 75 | 100 |
| --- | --- | --- | --- | --- |
| **Invertebrates** | 17 | 33 | 50 | 66 |
| **Fish** | 8 | 17 | 25 | 34 |

## SI 5B

### Summary of simulations

| **Initial Animals** | 25 | 50 | 75 | 100 |
| --- | --- | --- | --- | --- |
| **Food webs** | 50 | 50 | 50 | 50 |
| **Scenarios** | 2 | 2 | 2 | 2 |

## SI 6

### Differences between a small and large fishery

|  | Small fishery | Large fishery |
| --- | --- | --- |
| Net size (m) | 3.0 | 15.0 |
| Vessel speed (m^2^*s^-1^) | 1.5 | 1.5 (unchanged due to increased drag from larger net) |
| Maximum catch (g*s^-1^) | 23.0 (Ayunda et al., 2018)  listed as 5 gross tonnes capacity, with fishing trips 2.5 days = 23 g/s | 200.0 |
| fraction of day spent on fishing activities | 0.58  (ILO, 2004) 14 hours/24 hours | 0.9 |
| fraction of day spent actively fishing (ie not travelling to/from fishing areas) | 0.5 | 0.9 |

## SI 7

### Indicator descriptions

| Extrinsic Characteristics | |
| --- | --- |
| **Trophic similarity** | We used a Jaccard index to quantify how similar two species are in shared predator and prey species (0 indicates no similarity, 1 indicates complete niche overlap)(See Gauzens et al., 2015). |
| **Link Distance**: | The number of network links between the harvested and non-harvested species. |
| **Modules** | Modules are partitions of the food web that group together species that have more interactions between themselves than with species from other groups (Newman and Girvan, 2004). Our study differentiates between species belonging to the same module as the harvested species or to others. |
| **Role** | The interaction type between a non-harvested species and the harvested species. It can be Other-below, Prey, Competitor, Predator, or Other-above. Predator or Prey are determined through the food web; Competitor is a species within a 1.5 trophic level range around the harvested species and has a Jaccard similarity of at least 0.5; remaining species are classified as Other-above or Other-below if the non-harvested species has a trophic position above or below the harvested species, respectively (**SI 9**) |
| Intrinsic Characteristics | |
| **Interaction complexity** | The number of predator, prey, or competitor species a species has. |
| **Trophic level** | The sum of the trophic level of a consumer species’ resources times the fraction of these resource species that constitute the consumer’s diet plus one (Benke, 2011). Basal species start with a trophic level of one:  $TLi = \frac{1}{n}\sum_{j\epsilon prey}^{n} TLj + 1$ |
| **Trophic position difference** | The difference between the harvested species’ trophic position and a non-harvested species. |

## SI 8

### Role Plot

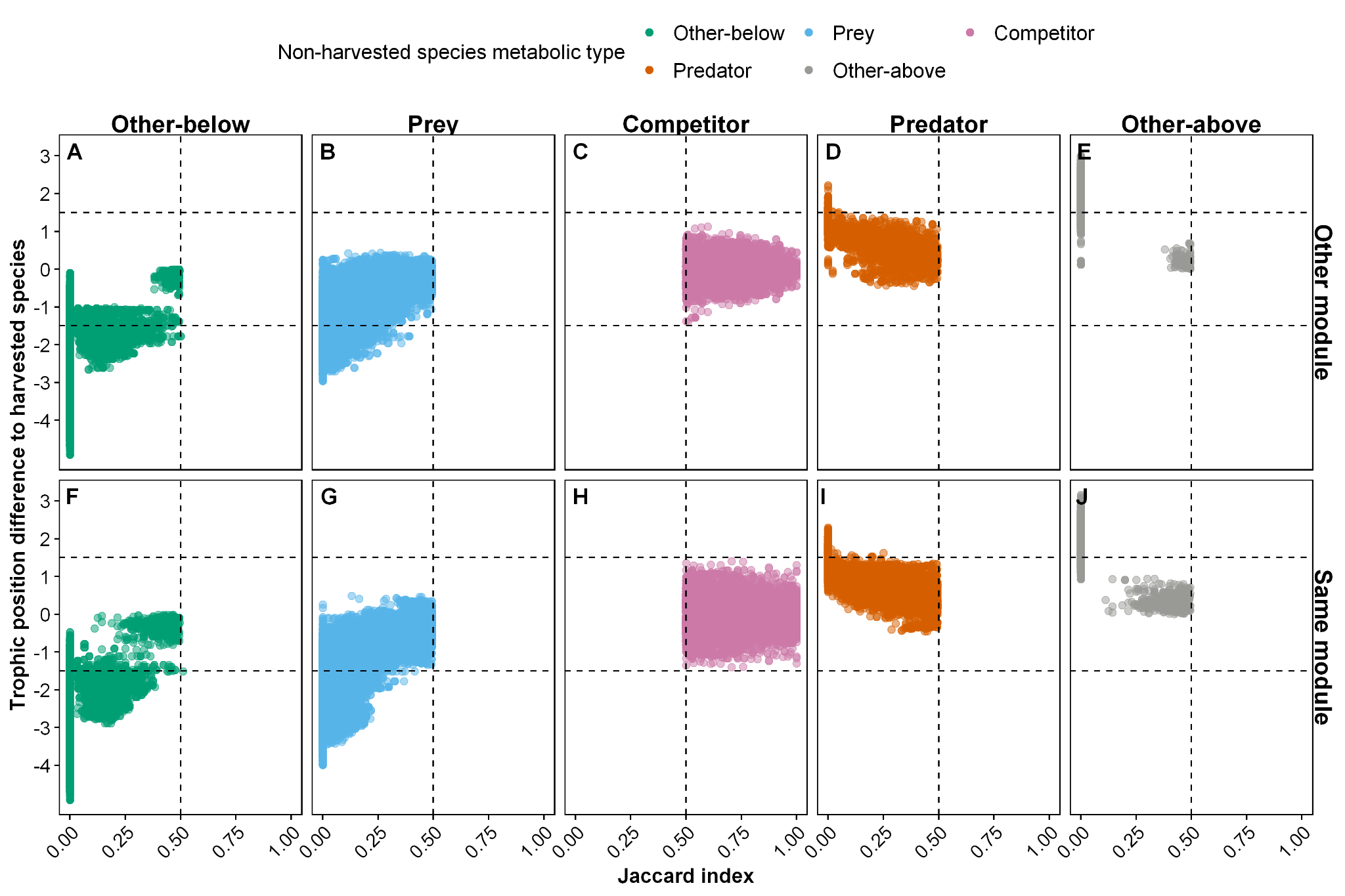

Assigning the species role for non-harvested species compared to their trophic position difference and Jaccard index to the harvested species. The Jaccard index (x-axis) shows the network connection overlap between a non-harvested species and the harvested species. The trophic position differenceto the harvested species (y-axis) shows the trophic position of the non-harvested species subtracted by the harvested species’ trophic position. The dashed horizontal lines at y = 1.5 and y = -1.5, and dashed vertical line at x = 0.5 are used to determine whether a species is a competitor of the harvested species. Each dot represents one species.

## SI 9

### Results of small fishery

*Direct Effects*

The best GAMs for harvested extinction (0.3 AIC difference for small, 2 for large) excludes metabolic type in both scenarios, while species richness was insignificant for the smaller fishery.

The best GAM model for the small fishery explained less variation in extinction (44% vs 73%) than the large fishery due to fewer extinctions occurring in the small fishery (0.8% vs 11%) or changes in an unknown variable.

*Indirect effects*

Although the mean relative biomass of a food web changes little (0.13 % and 0.39%, small and large), some non-harvested species achieved relative biomass increases of 272% and 5635% in the small and large fishery, respectively.

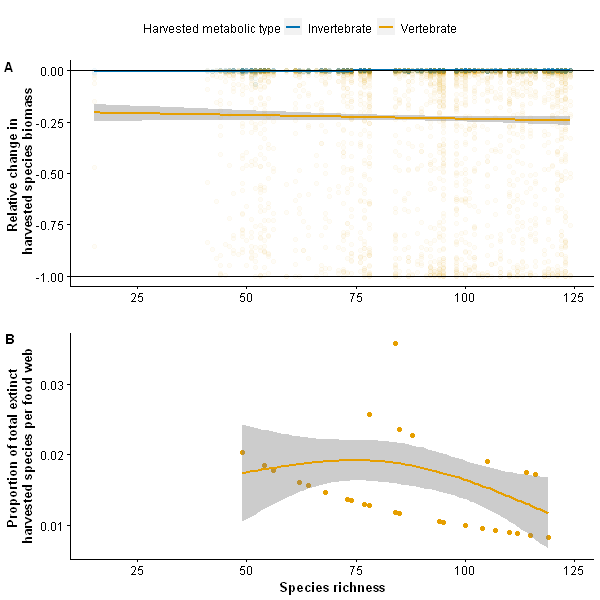

*Relative biomass change and extinction proportion of the harvested species for a small fishery. (A) Relative biomass change for harvested species (y-axis) vs food web species richness. Thin black lines indicate no change (y = 0) and species extinction (y = -1.0). Each faded point represents a harvested species. GAM with standard error for the relative change in harvested species biomass is calculated without extinct harvested species. (B) The proportion of total harvested extinctions per food web (y-axis) vs food web species richness. The proportion is the sum of all extinct harvested species that occurred in that food web divided by the number of species at the start of the simulation.*

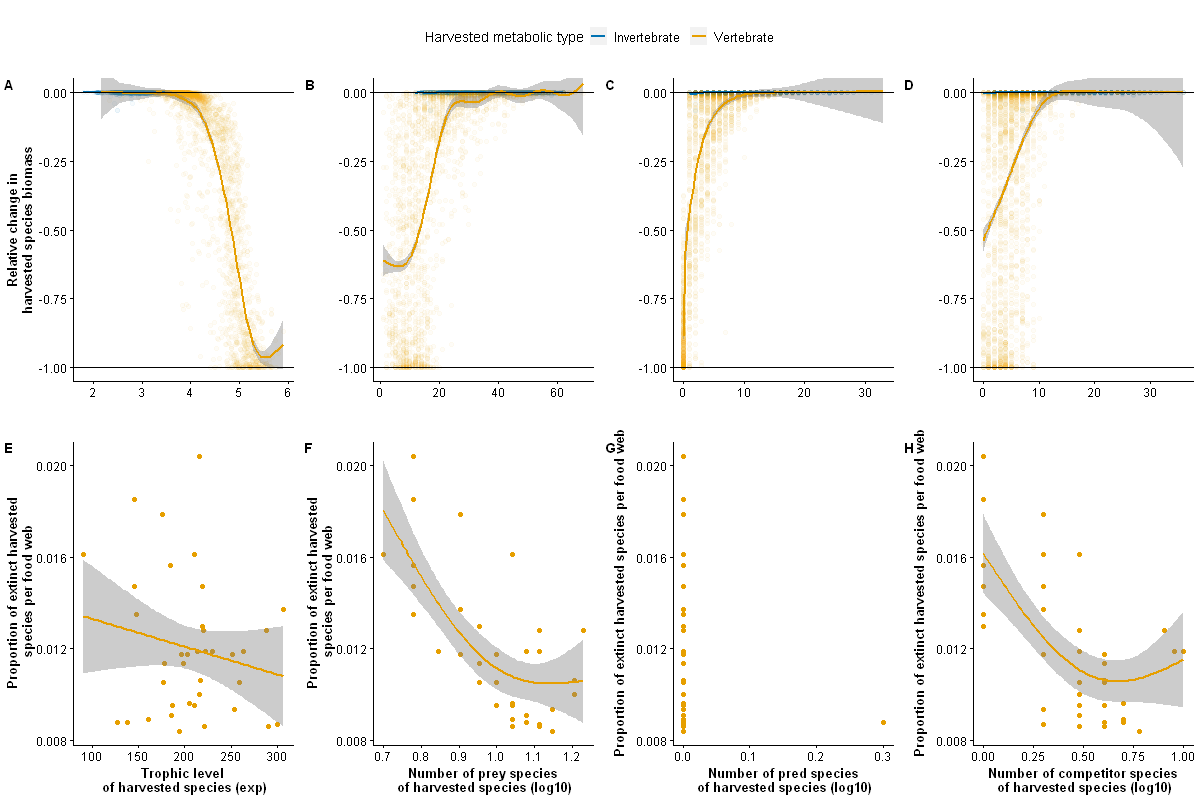

*Total extinction counts and biomass changes in the harvested species for a small fishery. Relative change in harvested species biomass (y-axis) vs the harvested species trophic position (A), number of prey species (B), number of predators (C), and number of competitor species (D) - Thin black lines indicate no change (y = 0) and species extinction (y = -1.0). Each faded point represents a harvested species. GAM with standard error for the relative change in harvested species biomass is calculated without extinct harvested species.*

*Proportion of harvested species extinction per food web richness (y-axis) vs the harvested species trophic position (E), number of prey species (F), number of predators (G), and number of competitor species (H). No invertebrate went extinct. The proportion was calculated by dividing the extinct harvested species by the number of species in that food web*

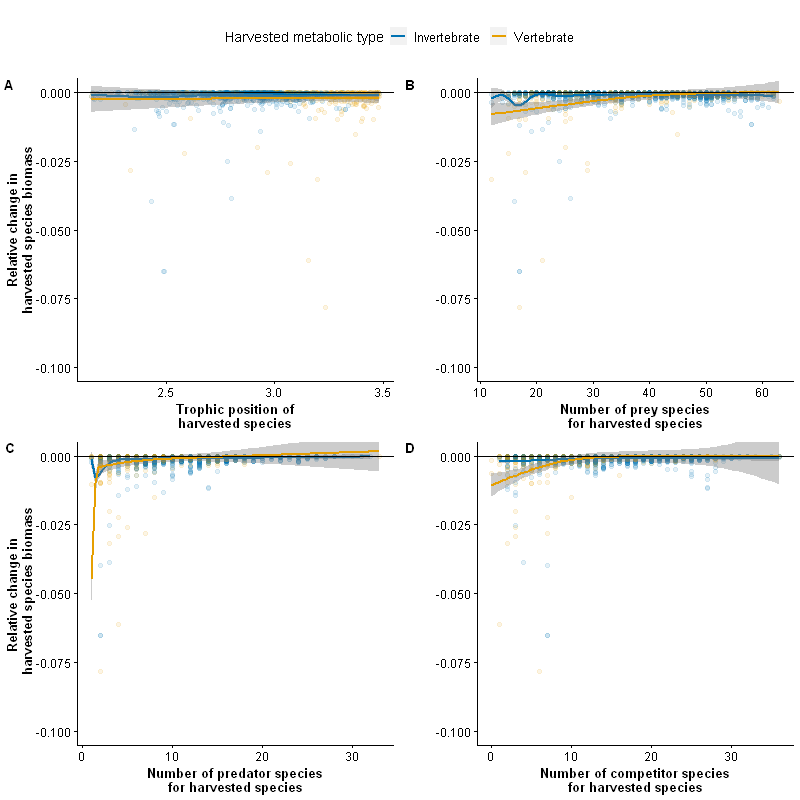

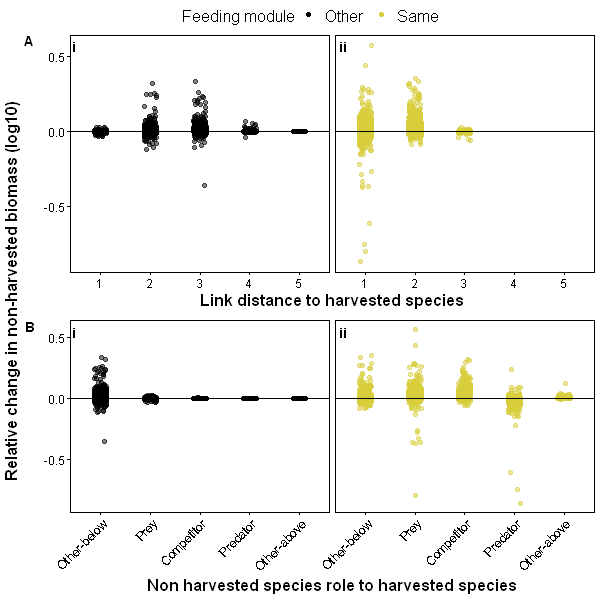

*Figure 2 Relative biomass change in non-harvested species for all food webs per link distance to, and role for, the harvested species, small fishery. The relative change in non-harvested species biomass (y-axis) is the harvested biomass divided by the pristine biomass for each non-harvested species. Link distance (x-axis) is the trophic position of the non-harvested species subtracted by the harvested species’ trophic position. Roles are assigned based on network connection and Jaccard Index. Species that interact closely are assigned to the same module. For each plot, the horizontal line indicates no change (y = 0). Each point represents a non-harvested species. Figure 2A Small Fishery results showing link distance and feeding module; Figure 2B Small Fishery results showing role and feeding module. Extinct species have been removed.*

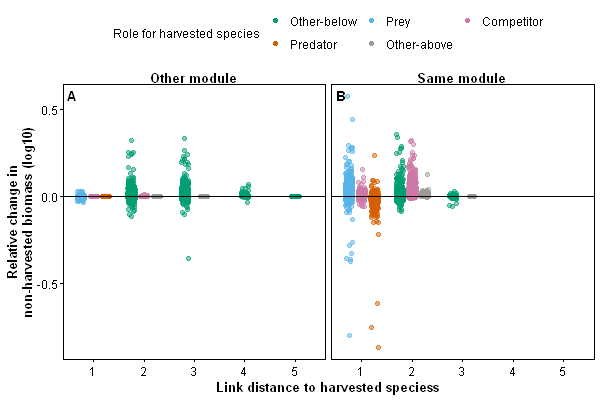

*Figure 3 Relative biomass change in non-harvested species per role and link distance to harvested species for all simulated food webs, small fishery. Relative change in harvested species biomass on a log10 scale (y-axis) versus the non-harvested species links distance to the harvested species (x-axis). Each point represents a non-harvested species. Thin black lines indicate no change (y = 0). Figure 3A Small Fishery results from a different feeding module; Figure 3B Small Fishery results from the same feeding module. Extinct species have been removed.*

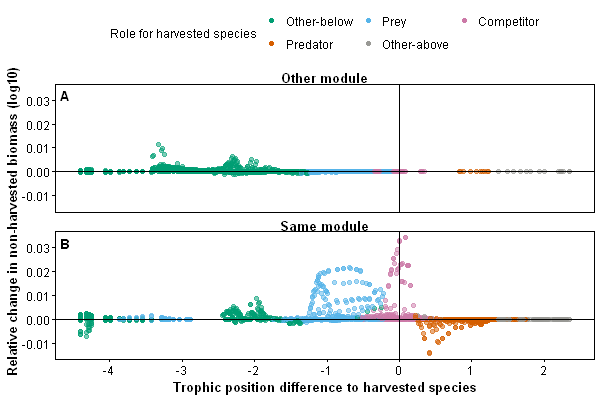

*Relative change in non-harvested biomass for a random food web, small fishery. The relative change in non-harvested species biomass (y-axis) is the harvested biomass divided by the pristine biomass for each non-harvested species. Trophic position difference (x-axis) is the trophic position of the non-harvested species subtracted by the harvested species’ trophic position. For each plot, the horizontal line indicates no change (y = 0), the vertical lines show trophic position differences of -3,-2,-1,0,1, and 2, with each point representing a non-harvested species. Figure 4A Small Fishery results from a different feeding module; Figure 4B Small Fishery results from the same feeding module.*

## SI 10

### Shared trophic levels for invertebrates and vertebrates (Large)

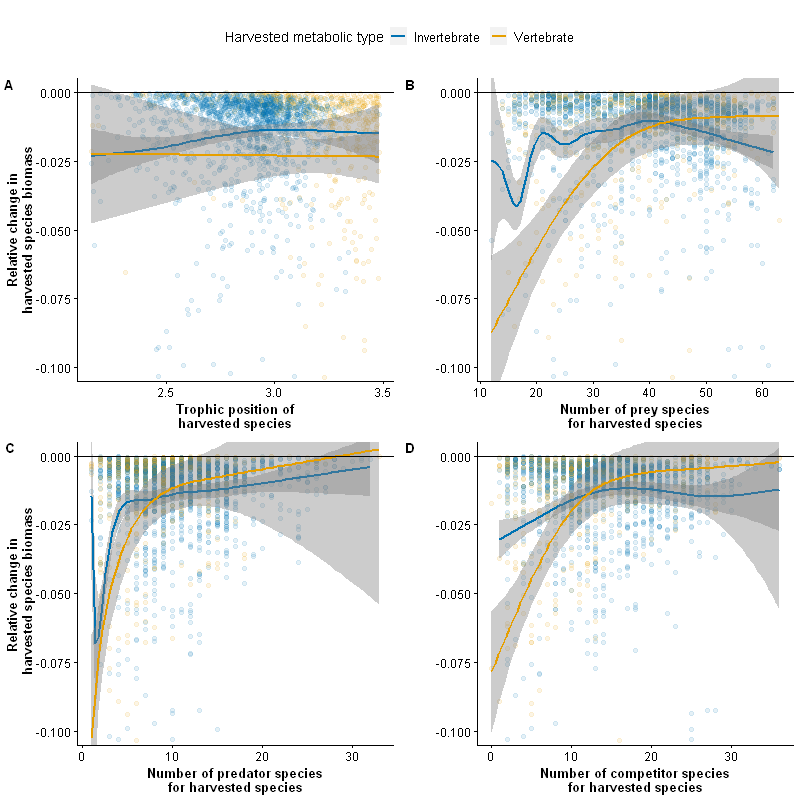
